# Creating a thermostable beta-glucuronidase switch for homogeneous immunoassay by disruption of conserved salt bridges at diagonal interfaces

**DOI:** 10.1101/2022.05.19.492583

**Authors:** Bo Zhu, Cheng Qian, Haoxuan Tang, Tetsuya Kitaguchi, Hiroshi Ueda

## Abstract

The *Escherichia coli* beta-glucuronidase (GUS) has been used as a reporter enzyme in molecular biology and has been engineered to enzyme switches for the development of homogeneous biosensors. Here, we developed a thermostable GUS enzyme switch from a thermostable GUS mutant TR3337 by disrupting a conserved salt bridge (H514-E523) between the diagonal subunits of its homotetramer. A combinatorial library (240 variants) was screened by a novel high-throughput strategy, and a mutant DLW (H514D/M516L/Y517W) was found to be a functional enzyme switch in a caffeine-recognizing immunosensor. The molecular dynamics simulations were performed to predict the topology change around position 514, and the sidechain flip of D514 (repulsion with E523) was observed in the DLW mutant. Up to 1.8-fold of the signal-to-background ratio was confirmed when measured at 45 °C, which makes the DLW mutant a versatile tool for developing the thermostable immunosensors for *in vitro* and *in cellulo* applications.

**Table of contents graphic:** 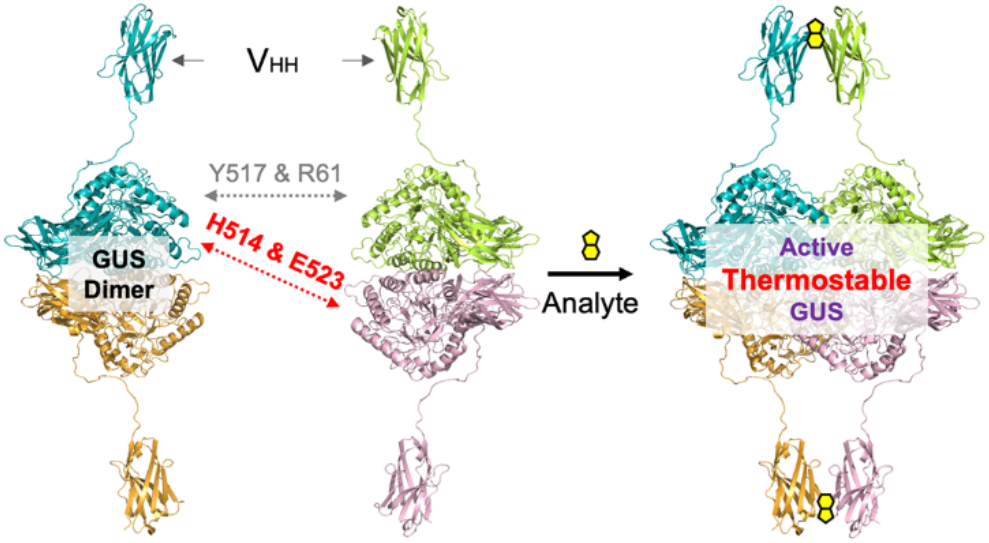

## INTRODUCTION

Homogeneous immunoassay attracts more attention in the field of point-of-care (POC) diagnostics under the background of the COVID-19 pandemic. Unlike the conventional heterogeneous immunoassay, e.g., Enzyme-Linked Immunosorbent Assay (ELISA), a homogeneous immunoassay is a one-pot system, which includes all the required components in bulk solution [1]. The simple operation, rapid readout, broad detection spectrum, and low cost make the homogeneous immunoassay also suitable for self-health monitoring, therapeutic drug monitoring, on-site drug abuse testing, food safety control, on-site pollution investigation, etc.

One major category of the homogeneous immunoassay functions through the property or distance change of fluorophores [2–5], nanoparticles [6], or fluorescence proteins [7] upon the binding of the analyte by the antibody part of the immunosensor (the analytes can also be antibodies). For example, the generalizable Quenchbody immunosensors are based on the fluorophore-quenching by intrinsic Trp of antibodies through photoinduced electron transfer and the dequenching upon analyte recognition [2]. Another major type of homogeneous immunoassay is based on the complementation [8–14] or allosteric regulation [15,16] of reporter enzymes upon the binding of the analytes.

The *E. coli* beta-glucuronidase (GUS) has been used as a reporter enzyme for over three decades [17,18] because of its stability and various commercially available substrates in fluorometric, colorimetric, and luminometric assays. The structure of the GUS was solved for its inhibitor study [19], and the tetramerization kinetics analysis of the GUS via *in vitro* translation revealed that the assembly of its active homotetramer occurs after the dimerization [20]. The mutations of two residues on the short interface (M516 and Y517) of the GUS were screened inside *E. coli* for creating a GUS mutant (M516<u>K</u>/Y517<u>E</u>, GUS_WT_-KE) with weakened tetramerization, which could be rescued by antibody-triggered complementation [8]. The GUS_WT_-KE has been used for generating homogeneous immunosensor for small molecules through fusing with variable domains (V_H_ or V_L_) of antibodies [9]. The affinity of V_H_ and V_L_ increases in the presence of antigen (known as open-sandwich principle [21]), which enhanced the tetramerization of the V_H_-GUS_WT_-KE and V_L_-GUS_WT_-KE dimers. A GUS mutant <u>IV-5</u> (GUS_IV5_) with a better thermostability [22] has been introduced into the above immunosensor to improve the stability and signal-to-background ratio (S/B) of this homogeneous immunoassay [10]. Four more mutation combinations at the 516 and 517 residues derived from the GUS_WT_-KE selection work were tested for finding a better interface mutant, GUS_IV5_-KW (M516<u>K</u>/F517<u>W</u>), with a lower background. The thermostability of the GUS_IV5_-KW-based immunosensor was improved compared with the original GUS_WT_-KE-based one. However, a 10-min incubation at 37 °C of one of the binary immunosensors, V_L_-GUS_IV5_-KW,still decreased the activity of GUS to around 60%. This will reduce the robustness of the GUS-based immunosensor while working at physiological conditions. Since the optimal temperature of the GUS_WT_ and GUS_IV5_ is 45 °C [23] and 65 °C [22], respectively, the significant negative effect of the interface mutations on the thermostability of GUS has been confirmed [10].

In this study, we used a hyperthermostable GUS mutant <u>TR3</u>337 (GUS_TR3_) [23] for further improving the thermostability of the GUS complementation-based homogeneous immunosensor at physiological temperatures. It was reported that the *E. coli* overexpressing GUS_TR3_ retained GUS activity after being treated at 100 °C for 30 min. Also due to its stability, the interface mutations identified in previous studies [8–10] could not break the tetramerization efficiently. Therefore, we identified a conserved salt bridge interaction (H514-E523_counter subunit_) between the diagonal subunits of GUS for the first time and found that additional alanine substitution of the H514 made the GUS_TR3_-KW a functional switch. To understand the fitness of the mutation combinations at 514, 516, and 517 residues, a combinatorial library (240 variants) was screened by a novel high-throughput strategy, and a mutant DLW (H514D/M516L/Y517W) showed the best switch function in the single-domain antibody V_HH_-fused immunosensor. Up to 1.8-fold of the signal-to-background ratio was confirmed when measured at 45 °C. The molecular dynamics (MD) simulations were performed to predict the topology change on the loop region around 514th residue, and the clear sidechain flip of D514 was confirmed in the GUS_TR3_-DLW mutant. The new mutant GUS_TR3_-DLW has a wider stable working temperature range than the previously reported GUS_IV5_-KW, which makes it a versatile enzyme switch for developing the thermostable homogeneous immunosensors for both *in vitro* and *in cellulo* applications.

## MATERIALS AND METHODS

### Materials

The *E. coli* strain XL10-Gold was purchased from Agilent for gene cloning. The *E. coli* SHuffle T7 Express lysY and the restriction enzymes were purchased from NEB Japan. The DNA polymerase (KOD-Plus-Neo) was purchased from Toyobo Biochemicals. Other chemicals were purchased from FUJIFILM Wako Pure Chemicals unless otherwise indicated. The oligonucleotides used in this study were synthesized by Eurofins Japan and listed in **Table S1**.

### Expression and Purification of Immunosensors

The expression plasmid was transformed into *E. coli* SHuffle T7 Express lysY. A 200 mL of LB medium (Difco LB broth, Lennox, BD) containing 100 μg/mL ampicillin was inoculated with 2 mL of overnight pre-culture. Cells were grown at 30 °C to OD_600_ ~ 0.8, then the expression of the immunosensor was induced with 0.5 mM isopropyl β-D-thiogalactopyranoside (IPTG) for 16 h at 16 °C with shaking at a speed of 200 rpm. The cells were harvested by centrifugation at 6,500 × *g* for 15 min at 4 °C. Then the pellet was suspended in 5 mL of lysis buffer (50mM sodium phosphate, 500 mM NaCl, pH7.4). The cells were disrupted by using an ultrasonicator (Qsonica Q125) in an ice-water mixture for 20 minutes (ultrasonication time) with 40% amplitude and 1 s interval. The lysate was recovered by centrifugation at 8,000 × *g* for 10 min at 4 °C. The supernatant was mixed with equilibrated TALON Metal Affinity Resin (200 μL of slurry, Takara Bio) for 1 h at 4 °C. The resin was then packed in a TALON 2 mL Disposable Gravity Column (Takara Bio), and the column was washed three times with 3 mL of washing buffer (50 mM sodium phosphate, 500 mM NaCl, 20 mM imidazole, pH 7.4). Proteins bound to the resin were eluted with 600 μL of elution buffer (25 mM sodium phosphate, 250 mM NaCl, 500 mM imidazole, pH 7.4) by incubation on ice for more than 20 min. The glycerol was added into the eluted protein to a final concentration of 50% and the protein solution was stored at −80 °C. The purified proteins were analyzed by SDS-PAGE on a precast gradient gel (5% - 12% SuperSep Ace).

### Homogeneous Immunoassay for Caffeine Detection

A 25 μL of the 400 nM immunosensor protein was first mixed with 25 μL of 40 μM caffeine (Nacalai Tesque, Kyoto, Japan) in phosphate-buffered saline (PBS) containing 0.1% Tween 20 (PBST). The mixture was incubated at 25 °C for 30 min, then transferred into a 96-well black microplate (675076, Greiner Bio-One), followed by adding 50 μL of 0.3 mg/mL substrate 4-methylumbelliferyl β-D-glucuronide (4-MUG) in PBST. The fluorescence intensity was measured by using a microplate reader (CLARIOstar, BMG Labtech). Triplicate measurement was taken with an interval of 1 min at 25 °C with an excitation wavelength of 360/20 nm and an emission wavelength of 450/30 nm at gain 300.

### Diagonal Interface Residues Analysis

The PyMOL (version 2.4.2) and InterfaceResidues script (from PyMOL Wiki, Jason Vertrees, 2009) were used for identifying the interactions between the residues at the diagonal interfaces of GUS. As the preparation step, the crystal structure of wild-type GUS (PDB: 3LPF) was imported into the PyMOL with assembly 1, and the statuses were split to generate two object molecules containing two subunits of each. The four subunits were renamed with different chain numbers and merged into one object. The interface residues at the diagonal interface were listed via the InterfaceResidues script with the dASA (changes in accessible surface area) cutoff set at 10. Then, the residues with the interactions (Pi-interactions and polar interactions) against the residues at the diagonal counter subunit were analyzed by PyMOL ‘find’ function.

### Combinatorial Library Construction

The library DNA fragments containing random mutations at H514, K516, and W517 positions were prepared by PCR from the template pET32-VHH-GS-GUS-TR3337_KW-FLAG with the mixed primer BZ-TR3337-Lib1-Ins-F, BZ-TR3337-lib1Y-R, BZ-TR3337-lib1W-R, and BZ-TR3337-lib1E-R at a molar ratio of 1:0.33:0.33:0.33. The vector fragment was obtained by an inverse PCR from the template pET32-VHH-GS-GUS-TR3337_KW-FLAG with primer BZ-TR3337-Lib1-Vec-F and primer BZ-TR3337-Lib1-Vec-R. The library DNA fragments and the vector fragment were ligated by using In-Fusion assembly and transformed into XL10-Gold competent cells. Over 10,000 transformants were pooled and cultivated for extracting the library plasmids.

### High-throughput Screening

The transformed *E. coli* SHuffle T7 Express lysY carrying the V_HH_-GUS_TR3_ mutants were inoculated into a 96-well microplate (V96 PP 0.45 mL, Nunc) with 200 μL of LB medium containing 100 μg/mL ampicillin (LBA) per well. The pre-culture plates were cultured overnight at 30 °C at 1,000 rpm. Ten microliters of the pre-cultures were inoculated into the main-culture plates with 200 μL of LBA medium per well followed by 2 h cultivation at 30 °C at 1,000 rpm. The overexpression was induced by adding 0.5 mM IPTG to each well, and the cultivation was continued at 16 °C for 16 h at 1000 rpm. After the induction, the *E. coli* cells were pelleted by centrifuging at 1,750 × *g* for 15 min, and resuspended in 100 μL of PBS. The cells were disrupted using an ultrasonicator (VCX-130PB, SONICS, Sonics & Materials) with 20% amplitude for 15 times (1 s interval) on ice. The microplates were centrifuged again, and 80 μL of supernatant of each well were equally split into two wells on the assay microplates with preloaded 10 μL of PBST or caffeine solution (100 μM in PBST). The assay plates were incubated at room temperature for 15–20 min, and 50 μL of 4-MUG substrate solution (0.3 mg/mL in PBST) were added to each well. The fluorescence was measured using a microplate reader as described above. The S/B ratio and the relative reaction rate were used for the evaluation of the picked clones.

For the precise evaluation of the selected clones, small-scale cultivation of the clones was performed to check the performance of the immunosensors in triplicate. A 4 mL of LBA medium was inoculated with 40 μL of overnight pre-culture. After the cells were grown to an OD_600_ of 0.7– 0.8, 0.5 mM IPTG was added for the induction at 16°C for 16 h. A 600 μL of the culture was pelleted and resuspended in 300 μL of PBS. The cells were then disrupted in a water-ice mixture by using an ultrasonicator (Qsonica Q125) with 30% amplitude 25 times (1 s interval). The supernatant was used for the evaluation of the sensing function and reaction rate of the selected clones as described above.

### Thermostability Profiling and Antigen Dose-Dependency Measurement

For the thermostability profiling, a mixture containing 100 nM immunosensor protein, 100 μM caffeine, and 0.15 mg/mL 4-MUG was prepared in PBST with 5% PEG 6000 on ice before the measurement. The assay mixture was transferred into a 96-well black microplate, and the fluorescence intensity was measured immediately by using a microplate reader with a pre-heated incubation chamber at 25 °C, 37 °C, or 45 °C. For analyte dose-dependency measurement, the temperature was set to 25°C, and triplicate measurement was performed with an interval of 1 min for an hour as described above. The reaction rate between 20–30 min incubation was estimated by linear fitting using Microsoft Office Excel. The blank sample (no analyte) was used for the normalization of the reaction rate. Dose-response curves were fitted to a four-parameter logistic equation (1) using SciDAVis software (version 2.4.0) while ‘a’ was set to 1 as a constant parameter. The limit of detection (LOD) was calculated as the concentration corresponding to the mean blank value plus its three times the standard deviation.

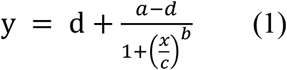

### MD Simulation

The GUS_TR3_ and GUS_TR3_-DLW PDB files were prepared from the wild-type GUS structure (PDB: 3LPG) with mutagenesis wizard in PyMOL after molecule preparation described in the interface analysis section (all selenomethionine residues were replaced with methionine and the inhibitor molecules were removed). The MD simulations were performed using the GROMACS 2021 [24]. The topology files were generated from the prepared PDB files while AMBER99SB force field [25] and default SPC/E water model were used. The systems were then solvated with the SPC216 water model and neutralized with sodium ions in cubic boxes with a volume of 4539 nm^3^. The short-range Van der Waals and electrostatic cut-off were set to 1.0 nm through all the simulations. The periodic boundary conditions were applied to all axes. The steepest descent minimization algorithm was used for energy minimization until the maximum force is lower than 1000 kJ/mol/nm. The NVT and NPT equilibrations were performed while the LINCS algorithm was used for holonomic constraints of H-bonds, and the particle mesh Ewald method was used for dealing with long-range electrostatic interactions. The NVT equilibration was performed for 1 ns with the Berendsen thermostat algorithm at 298.15 K. The NPT equilibration was performed for another 1 ns with the Parrinello-Rahman algorithm at 1 bar in isotropic pressure coupling type. The MD simulation was performed for additional 15 ns for each system. The RMSD was analyzed with GROMACS gmx-toolbox and visualized with Python (v3.7.12) library matplotlib (v3.2.2). The RMSD of each system reached a stable phase after 10 ns simulation (Figure S1).

## RESULTS AND DISCUSSION

### Prototype of Caffeine Immunosensor with GUS_TR3_ as Switch Enzyme

To evaluate the feasibility of the hyperthermostable GUS_TR3_ [23] as the switch enzyme in a homogeneous immunosensor, the immunosensor construct V_HH_-GUS_IV5_-KW (M516<u>K</u>/F517<u>W</u>) for caffeine detection [10] is utilized as a model by replacing the GUS_IV5_ with GUS_TR3_. The single-domain anti-caffeine antibody V_HH_ (clone VSA2) can be dimerized in the presence of its antigen caffeine [26], which triggers the tetramerization of the GUS dimers (**Figure 1A**). This V_HH_ is a thermostable antibody fragment, and it retained over 90% binding activity after 20 min incubation at 90 °C [27], which makes it a good model antibody for checking the immunosensor function at higher temperatures. As shown in **Figure 1B**, the previously reported GUS [9] or GUS_IV5_-based [10] enzyme switches were achieved by introducing mutation at positions 516 and 517 to weaken the subunit 1-3 and 2-4 interactions (**Figure 1A**). The same strategy was tested on the V_HH_-GUS_TR3_ caffeine immunosensor prototypes V_HH_-GUS_TR3_-KW (M516<u>K</u>/Y517<u>W</u>) and V_HH_-GUS_TR3_-KE(M516<u>K</u>/Y517<u>E</u>). The caffeine-dependent activity increase (sensing function) of both prototypes was negligible (**Figure 1C and 1D**). To eliminate the possibility that a misfolded V_HH_ caused the disfunction of the immunosensor, the ELISA was performed for confirming the binding between BSA-caffeine conjugate and V_HH_-GUS_TR3_-KE. A strong caffeine binding signal was observed at 100 nM concentration of the immunosensor (**Figure 1E**), which indicated that the small S/B was not due to the misfolding of the antibody region, but the insufficient separation of the GUS_TR3_ tetramer with current interface mutations. It has been reported that thermostable GUS_TR3_ has a strong tetramerization tendency [23], therefore, we started looking for other critical residues assisting the tetramerization to introduce additional mutations.

**Figure 1.**
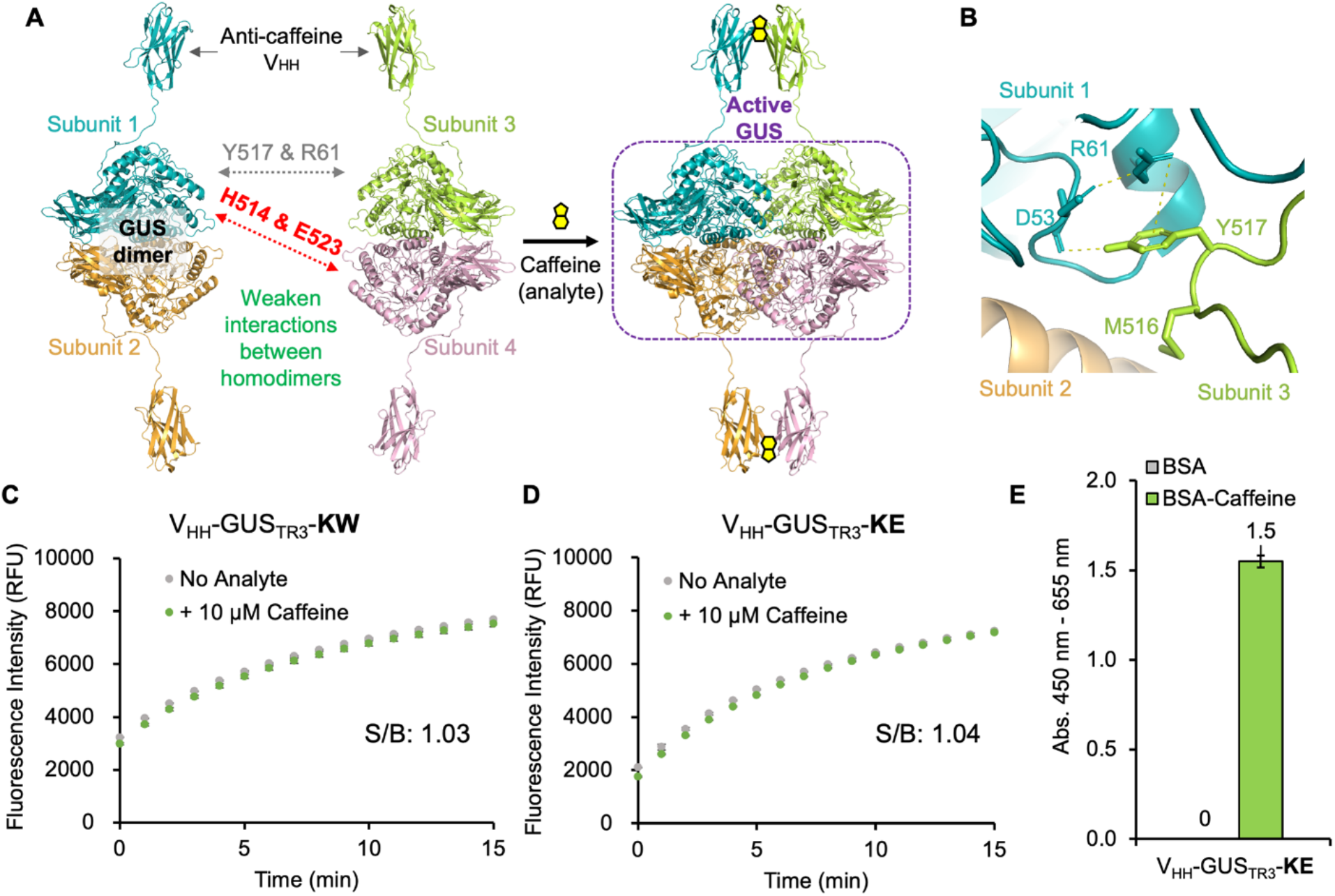
GUS complementation-based homogenous immunosensor. (A) Schematic illustration of the V_HH_-GUS immunosensor. The V_HH_(VSA2)-GUS_TR3_ monomer was modeled by AlphaFold and assembled via superimposing to the structure (PDB: 3LPG) of wild-type GUS. The linker region and the V_HH_ were adjusted manually in PyMOL. (B) Interchain interactions between the Y517 and the counter residues in wild-type GUS (PDB: 3LPF). (C) Time course activity measurement of V_HH_-GUS_TR3_-KW. (D) Time course activity measurement of V_HH_-GUS_TR3_-KE. S/B was calculated according to the reaction rate from 0–5 min. (E) ELISA result showing the binding between the V_HH_-GUS_TR3_-KE and caffeine. All data are expressed as mean ± standard deviation (n = 3).

### Discover Conserved Salt Bridges at Diagonal Interfaces of GUS

The mutations introduced at M516 and Y517 on the interface between the subunits 1-3 and 2-4 (**Figure 1A and 1B**) have been proved effective for weakening the dimerization of GUS [8,9] or GUS_IV5_ [10], but not sufficient for GUS_TR3_. Therefore, we decided to focus on looking for the residues involved in the interactions between diagonal interfaces if any. Firstly, all the residues close to the diagonal interface of GUS were listed using PyMOL (**Table S2**). Then, the polar interactions and Pi-interactions with other residues of the diagonal counter subunit were analyzed for each of the listed 13 residues. Finally, only one pair of salt bridge interactions was discovered between H514 and E523 as shown in **Figure 2A**. Notably, the M516 and Y517 are also close to the diagonal interface but do not have interaction against the diagonal subunit. To understand that if this pair of salt bridge is conserved during the evolution, the amino acid sequence of GUS was aligned with the GUS sequence or GUS-like sequence from other organisms with an identity greater than 40% (**Table S3**). The sequences from A512 to Y525 were extracted and converted to a WebLogo in **Figure 2B**. We found the H514-E523 motif is quite conserved across the GUS or GUS-like enzyme from different organisms. In contrast, the M516 and Y517 only exist in part of the bacterial GUS. Therefore, we decided to break the H514-E523 interactions by alanine substitution to see if it will weaken the tetramerization of GUS_TR3_. Since the E523 is also involved in other hydrogen bonds formation within the subunit (**Figure 2C**) and is part of a helix, the H514 was selected for introducing the mutation. The V_HH_-GUS_TR3_-AKW (H514<u>A</u>/M516<u>K</u>/Y517<u>W</u>) and V_HH_-GUS_TR3_-AKE (H514A/M516<u>K</u>/Y517<u>E</u>) were constructed and their immunosensor functions were evaluated (**Figure 2D and 2E**). Interestingly, the V_HH_-GUS_TR3_-AKW showed a 1.14-fold signal increase in the presence of 10 μM caffeine, however, the GUS activity of the V_HH_-GUS_TR3_-AKE was completely diminished. The SDS-PAGE of the purified immunosensor proteins indicated that their expression or solubility was not affected by the mutation significantly (**Figure 2F**). Therefore, we hypothesized that the mutation combination at the 514/516/517 residues is critical for achieving a GUS_TR3_ mutant with improved enzyme switch function.

**Figure 2.**
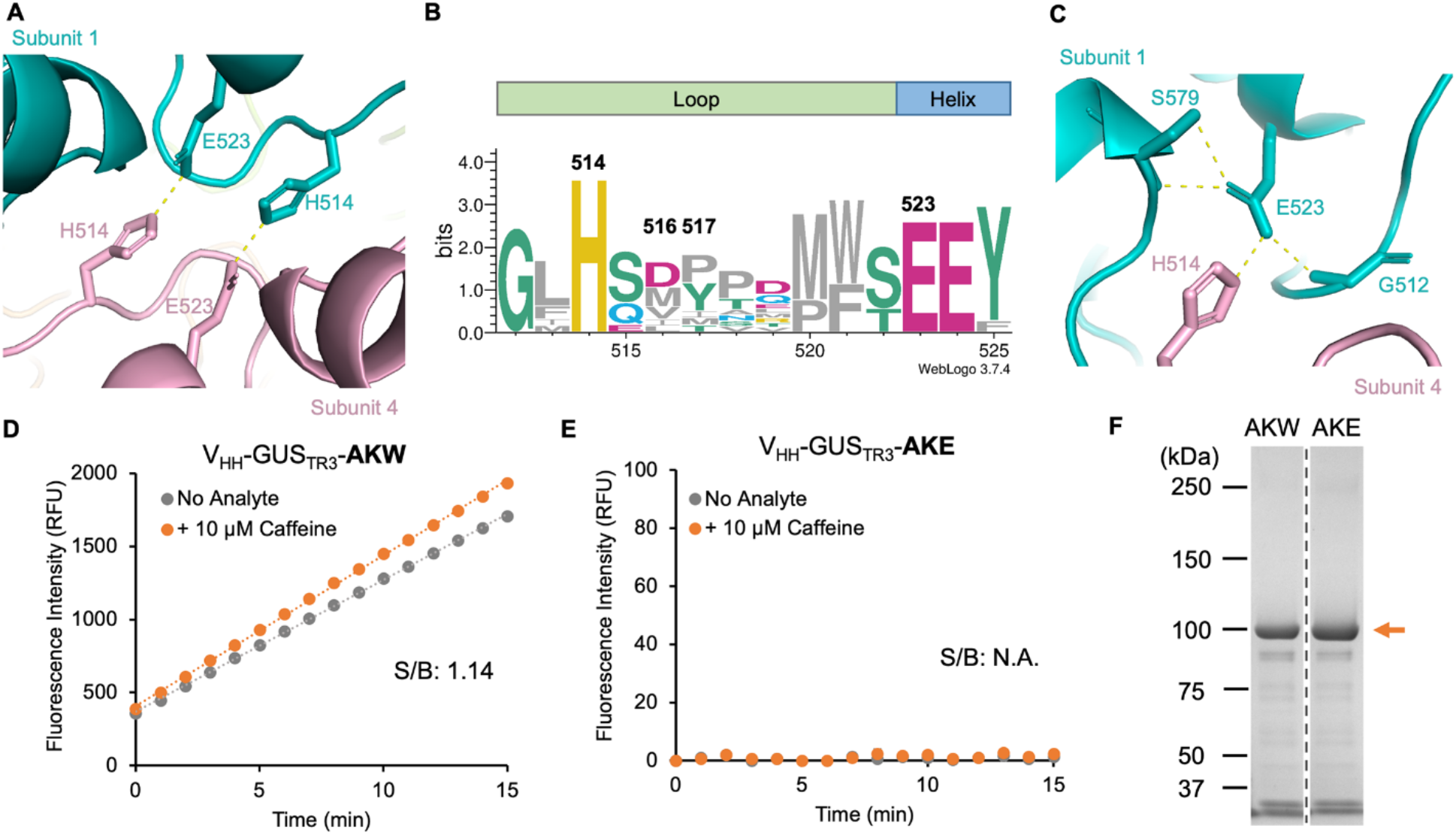
Conserved salt bridges at diagonal interfaces of GUS and the effect of Ala substitution on the switch function. (A) H514-E523 salt bridge between the diagonal subunits of wild-type GUS (PDB: 3LPF). (B) WebLogo (residue 512–525) showing the alignment of GUS or GUS-like scaffold from eleven organisms with amino acid identity > 40%. Color scheme of amino acid: polar, green; neutral, blue; basic, yellow; acidic, purple; hydrophobic, grey. (C) Residues show interactions with E523 (PDB: 3LPF). (D) Time course activity measurement of V_HH_-GUS_TR3_-AKW. S/B was calculated according to the reaction rate from 0–15 min. (E) Time course activity measurement of V_HH_-GUS_TR3_-AKE. All data are expressed as mean ± standard deviation (n = 3). N.A.: Not available. (F) SDS-PAGE analysis of the purified immunosensor variants. AKW: V_HH_-GUS_TR3_-AKW; AKE: V_HH_-GUS_TR3_-AKE. The orange arrow indicates the monomer of the immunosensors. Dash line divides the grouped lanes that are from two gels.

### High-throughput Screening of a Combinatorial Library of V_HH_-GUS_TR3_

To find the better mutation combination at the 514/516/517 residues of GUS_TR3_, we decided to perform a screening of a combinatorial immunosensor library. Previously, a glutathione S-transferase (GST) dimerization-based screening method has been reported for the selection of the ideal GUS mutants with a switch function [8]. The selection was performed according to the GUS activity of *E. coli* colonies hydrolyzing a colorimetric substrate on LB plates. The GUS mutants did not show activity without fusing with GST, but restored activity after fusing with GST, were considered as the functional clones. The gene of GUS mutants was required to be subcloned between two plasmids, which could introduce biases and difficulties for automation. Therefore, we established a new high-throughput screening method for the selection of functional GUS enzyme switch mutants based on the caffeine homogenous immunosensor design (**Figure 3A**). The *E. coli* transformants carrying the V_HH_-GUS_TR3_ mutants were inoculated into a 96-well microplate and cultured overnight as the pre-culture plate. Part of the pre-cultures was inoculated into the main-culture plate followed by another overnight induction. This two-step cultivation mimicked the GUS-based immunosensor overexpression process and this method would reduce the bias caused by the colony picking and growth speed variation among clones. After the induction, the *E. coli* cells were pelleted and resuspended in PBS buffer for cell lysis using an ultra-sonicator. Next, the microplates were centrifuged again, and the supernatant of each well was equally split into two wells on assay microplates with or without pre-loaded caffeine solution. The assay plates were incubated at room temperature to trigger the GUS tetramerization, and the fluorogenic substrate solution was added right before the fluorescence measurement using a microplate reader. The signal-to-background ratio (S/B) and the relative reaction rate were used for the evaluation of the picked clones. For the preparation of the combinatorial library (diversity = 240, excluding variants with stop codon), saturated mutagenesis was introduced on the H514 residues with NNS degenerate codon for a better following of the frequency of codon usage of *E. coli* tRNAs [28]. For position 516, M (parental), K, Q (two amino acids selected in previous GUS switch work [8]), and L were introduced with the MWG degenerate codon. For position 517, Y (parental), W, and E (two amino acids selected in previous GUS switch work [8]) were introduced with three different codons. 283 valid clones (growth confirmed) were subjected to the screening process, and the results are shown in **Figure 3B**. The calculated probability of discovering at least one of the top 2 variants from 283 clones is over 98% [29]. The majority of the clones exhibited low S/B while the reaction rate showed a wide distribution. Interestingly, there was a trend that more activity-reduced clones showed better sensing function. This phenomenon indicated that there is a trade-off between the weakened tetramerization and the activity for the hyperthermostable GUS mutant TR3337. The clones above the criteria we chose were re-evaluated in a triplicate manner and sequenced, and the results of the single clones were shown in **Figure 3C**. We found two mutants, DLW and SLE, appeared twice in the selected clones, and the AKW tested above was also selected during our screening. The Control-AKE showed almost no activity, while Control-KW showed the highest activity but no sensing function when subjected to this screening, which was consistent with the result of the purified immunosensors. Therefore, we decided to choose DLW and SLE variants for further evaluation after purification. Since two QY (M516Q/Y517Y) variants, AQY and LQY, were selected, we decided to also test the AQY variant to compare with the previously tested AKW mutant. Notably, the QY (516/517) variant were also selected in the previous work for making the enzyme switch based on the wild-type GUS [8].

**Figure 3.**
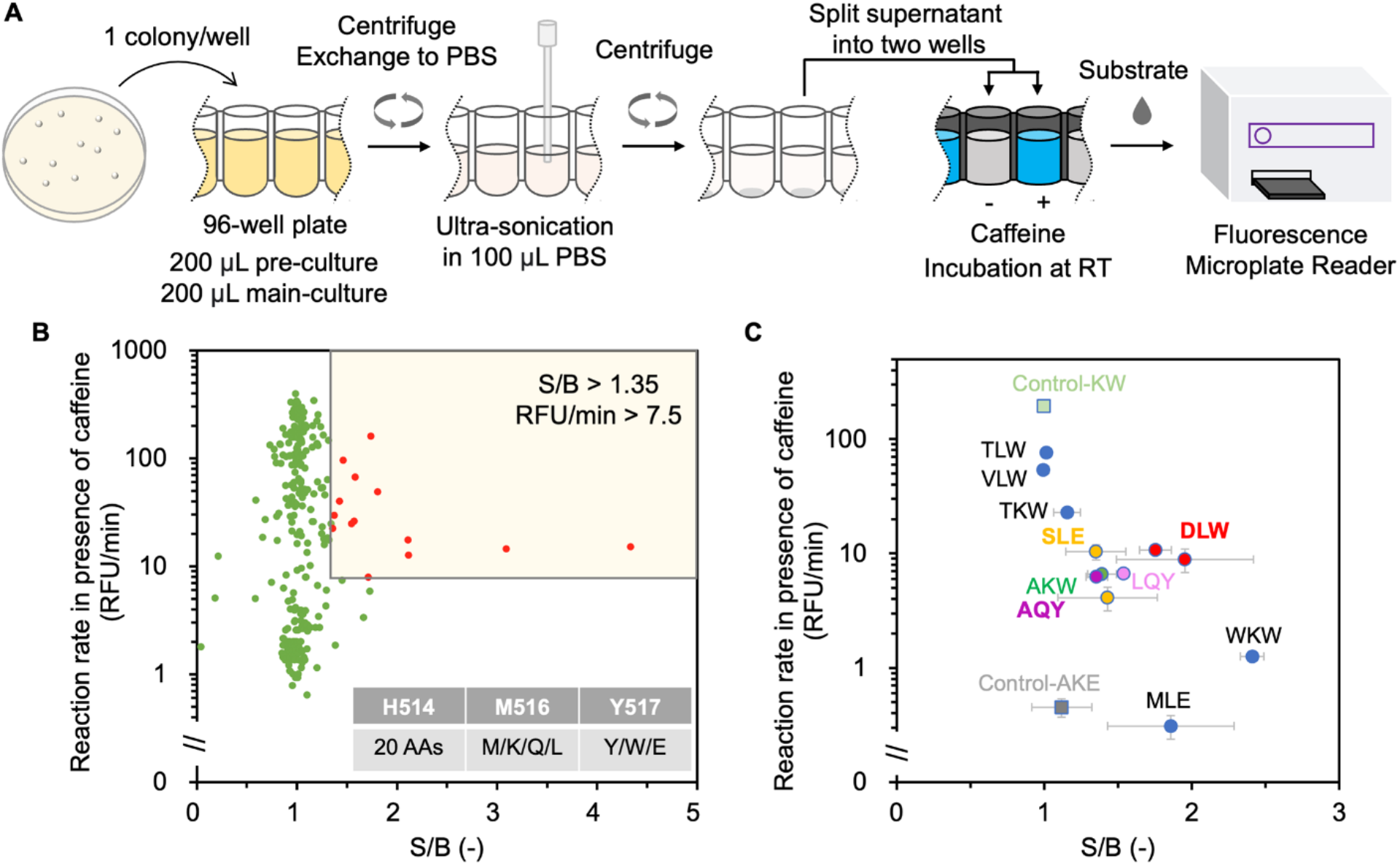
High-throughput screening of a combinatorial library of V_HH_-GUS_TR3_. (A) Schematic representation of the novel high-throughput screening method. RT: room temperature. (B) Screening results of 283 clones (n =1). S/B was calculated according to the fluorescence intensity increase from 0—15 min. Red dots represent the clones that went into the second-round triplicate evaluation. (C) Evaluation of the selected clones and the substitutions at residues 514/516/517 in every single clone. S/B was calculated according to the reaction rate from 0–14 min. Data are expressed as mean ± standard deviation (n = 3).

### Functional Analysis of Selected GUS_TR3_ Variants

Three caffeine-recognizing immunosensors carrying GUS_TR3_ mutants DLW, SLE, and AQY were expressed and purified (**Figure S2**) for evaluating their function under defined conditions. The purified DLW variant exhibited the best S/B (**Figure 4A**), which is correlated to the screening result (**Figure 3C**). The SLE variant showed a similar S/B to the AKW variant (**Figure 4B**), and it showed that the Y517E mutant-based variant would also be functional if an ideal combination of mutations is introduced at positions 514 and 516. The AQY variant (**Figure 4C**) showed better S/B than SLE and AKW, but was not able to compete with DLW. An obvious drop of the reaction rate was also confirmed within all selected mutants, especially for the DLW mutant. It should be noticed that the L at position 516 in the library was introduced because of the usage the MWG degenerate codon with no rationale. However, the L516 appeared several times in the selected variants including the best DLW mutant. It showed the possibility that the chosen M/K/Q at position 516 according to the previous work [8] was not the most preferred substitution for making the GUS_TR3_-based enzyme switch. Another screening of the DLW-based library with the saturated mutagenesis at position 516 and other positions (e.g. 513 or 523) could be performed in the future to gain a better understanding of the fitness of the mutation combination around position 514. **Figure 4D** shows the loop (in blue) connecting the catalytic residue E504 and the beginning of the helix (E523). This loop includes the three critical residues (514/516/517) for designing a functional enzyme switch from GUS. The change of the reaction rate of the selected mutants after the tetramerization could be due to the accumulated displacement of E504. To understand why the DLW mutant exhibited a better switch function from a structural level, the MD simulations for GUS_TR3_ and GUS_TR3_-DLW were performed. **Figure 4E** shows the simulated structure of the DLW variant at 15 ns, and the H514 residue of the GUS_TR3_ at 15 ns is also shown for comparison. A clear sidechain flip of the D514 (repulsion with E523) in DLW mutants was confirmed, while the H514 in GUS_TR3_ kept the orientation of the sidechain with a positive charge towards the E523 residue. The sidechain flip could be observed starting from the 5 ns of the simulation, which is possibly a result of the minus charge repulse between D514 and E523. This predicted topology changes around residue 514 could lead to the improved switch function of the DLW variant.

**Figure 4.**
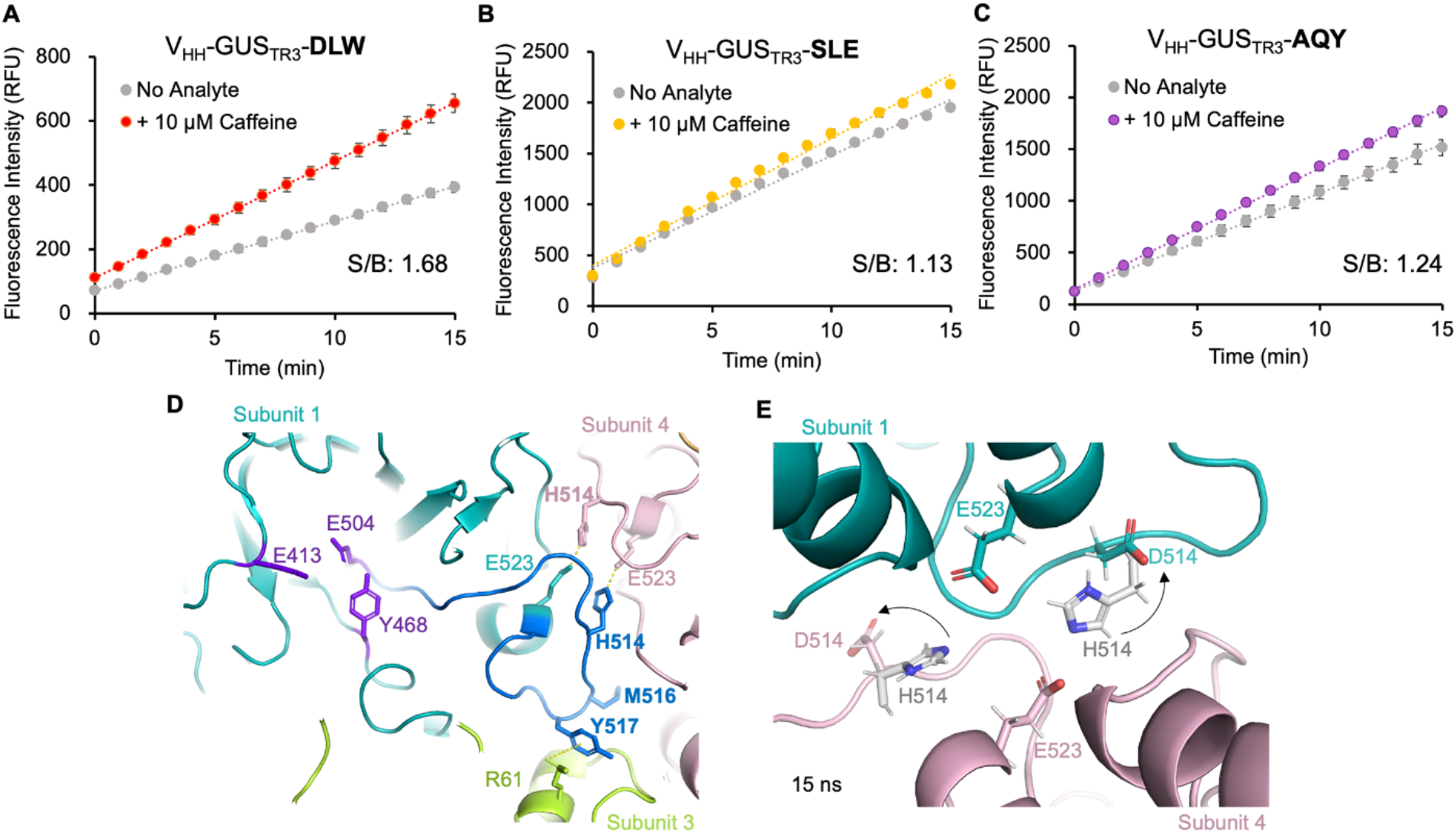
Functional and structural analysis of the immunosensors carrying the selected GUS_TR3_ mutants. (A) Time course activity measurement of V_HH_-GUS_TR3_-DLW. (B) Time course activity measurement of V_HH_-GUS_TR3_-SLE. (C) Time course activity measurement of V_HH_-GUS_TR3_-AQY.S/B was calculated according to the reaction rate from 0–15 min. All data are expressed as mean ± standard deviation (n = 3). (D) Detailed structure around the catalytic center (residues in purple) and interface mutations. The loop from catalytic residue E504 to E523 is labeled in dark blue. (E) MD simulation result of the GUS_TR3_-DLW at 15 ns. The simulated structure of GUS_TR3_ at 15 ns is superimposed and the H514 residues only are shown (grey) for comparison. The black arrow indicates the flip of the sidechain at residue 514. The O and N atoms in sidechains are labeled with red and blue, respectively.

### Improved Thermostability of V_HH_-GUS_TR3_-DLW Immunosensor

The thermostability is an important property to evaluate for the GUS_TR3_-based homogeneous immunosensor. Previously reported GUS_IV5_-based immunosensor lost 30–40% GUS activity after heating half of the binary immunosensors to 37 °C for 10 min [10]. Also, the sensing function of the immunosensors was not evaluated at higher temperatures. Therefore, in this study, we examined the sensing function of the selected GUS_TR3_-DLW-based immunosensor at 25 °C, 37 °C, and 45 °C to achieve a better understanding of its thermostability. To monitor the dynamics of the activity change, we started the fluorescence measurement immediately after mixing the immunosensor, analyte, and substrate at certain temperatures (**Figure 5A-C**). The increased reaction rate of the GUS_TR3_-DLW was confirmed at 37 °C and 45 °C, while the S/B did not change significantly. The reaction rates of different time ranges were analyzed to compare the activity dynamics at different temperatures (**Figure 5D**). At 25 °C, a stable reaction rate was achieved after 20 min incubation and no decrease was observed until 60 min. At 37 °C, the reaction rate reached a stable phase after 10 min incubation, and it was 3-fold faster than that of 25 °C between 10—20 min. At 45 °C, the reaction rate increased faster than the other two temperatures within 20 min, and a 4-fold reaction rate was achieved compared with that of 25 °C. After 20 min incubation at 45 °C, the reaction rate started decreasing and 50% of the activity retained after 60 min incubation. The thermostability profile of the immunosensor V_HH_-GUS_TR3_-DLW is completely different from the previously reported GUS_IV5_-KW-based one [10]. This result demonstrated that the novel GUS_TR3_-DLW enzyme switch was suitable for making the homogenous immunosensors that are designated to work under physiological or higher temperatures. The dose-dependency of the V_HH_-GUS_TR3_-DLW was also investigated at 25 °C, and the EC_50_ and LOD were calculated to be 2 μM and 107 nM, respectively (**Figure 5E**). Compared with the GUS_IV5_-KW-based immunosensor (EC_50_: 1.2 μM; LOD: 40 nM), the sensitivity is slightly decreased, which could be due to the relatively lower S/B of the V_HH_-GUS_TR3_-DLW immunosensor and the different affinity between the homodimers of the GUS mutants.

**Figure 5.**
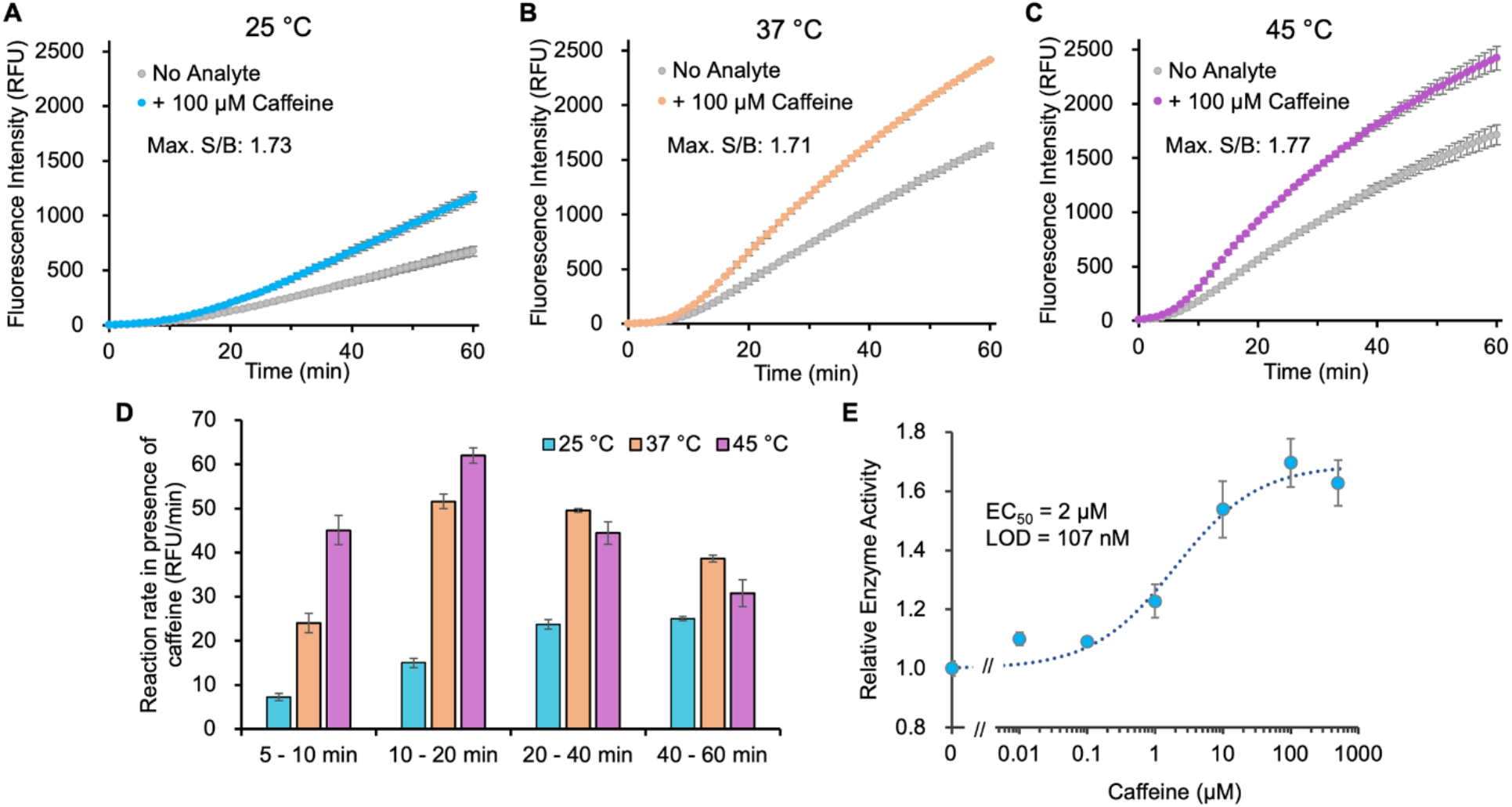
Thermostability profiling of V_HH_-GUS_TR3_-DLW immunosensor and its analyte dosedependency measurement. Time course activity measurement of V_HH_-GUS_TR3_-DLW at 25 °C (A), 37 °C (B), and 45 °C (C). S/B was calculated according to the fluorescence intensity at each time. Max.: maximum. (D) The reaction rate in the presence of 100 μM caffeine of different time ranges. (E) Caffeine dose-dependent curve. All data are expressed as mean ± standard deviation (n = 3).

## CONCLUSIONS

In this study, we identified a conserved salt bridge interaction (H514-E523) between the diagonal subunits of the GUS enzyme. A thermostable GUS enzyme switch, GUS_TR3_-DLW, was successfully developed by weakening the interactions between the homodimers via introducing the mutations at the H514, M516, and Y517. During this process, a high-throughput screening strategy was established and used for finding the optimal combination of the mutations at different residues. The MD simulation result of the DLW variant suggested the sidechain of the D514 flipped to the opposite direction of the parental H514, which could lead to its better switch function. The thermostability profile of the GUS_TR3_-DLW-based caffeine immunosensor was investigated and its function within and above physiological temperature was confirmed. This novel GUS enzyme switch, GUS_TR3_-DLW, has a broader stable working temperature than the previously reported GUS_IV5_-KW, which makes it a versatile tool for developing the thermostable homogeneous immunosensors and immunoassays for *in vitro* and *in vivo* applications. In addition, the screening method demonstrated here can be used for the development of other GUS mutants or immunosensors with an improved sensing function, specific activity, or thermostability.

## ASSOCIATED CONTENT

### Supporting Information

Supplementary methods; RMSD analysis results; SDS-PAGE for proteins; Oligonucleotides and synthesized gene sequences; Amino acid residues at diagonal interface of GUS; Alignment of the GUS sequences (PDF)

### Accession Codes

*E. coli* beta glucuronidase: UniProtKB P05804.

## AUTHOR INFORMATION

### Notes

The authors declare no conflicts of interest.

## ACKNOWLEDGMENTS

We thank the Biomaterials Analysis Division, Open Facility Center, Tokyo Institute of Technology for DNA sequence analysis. We also thank Dr. Jasmina Damnjanovic from Nagoya University for critical reading of the manuscript.

## ABBREVIATIONS

BSA: bovine serum albumin
EC_50_: half maximal effective concentration
GUS: *E. coli* betaglucuronidase
LOD: limit of detection
PBS: phosphate-buffered saline
RMSD: root mean square deviation
S/B: signal-to-background ratio
SDS-PAGE: sodium dodecyl sulfate-polyacrylamide gel electrophoresis

