## Supplementary Material for "Creating a thermostable beta-glucuronidase switch for homogeneous immunoassay by disruption of conserved salt bridges at diagonal interfaces"

Hiroshi Ueda

Laboratory for Chemistry and Life Science, Institute of Innovative Research, Tokyo Institute of Technology, Nagatsuta-cho, Midori-ku, Yokohama 226-8503, Japan


### Supplementary Methods

#### Plasmid constructions

The C-terminal sequence of the $GUS_{TR3}$ (from residue W492 to Q604) was synthesized by Integrated DNA Technologies (**Table S1**), and amplified by using polymerase chain reaction (PCR) with the primer Q493R_top and the primer GUS_XhoFor. The remaining sequence of GUS fragment (from residue M1 to A491) was amplified from the template pCA24N-GUS [1] by PCR with the primer GUS_NotBack and the primer Q493R_bottom. To connect these two $GUS_{TR3}$ fragments, overlap extension PCR was performed using the primer GUS_NotBack and the primer GUS_XhoFor. The GUS-TR3337_KE fragment was digested by the restriction enzyme NotI and XhoI, and was inserted into the linearized pET32-VHH(Caf)-GS-GUS-IV5_KW plasmid [2] (digested by the same restriction enzymes) to yield the pET32-VHH(Caf)-GS-GUS-TR3337_KE plasmid. The pET32-VHH(Caf)-GS-GUS-TR3337_KW plasmid was prepared from pET32-VHH(Caf)-GS-GUS-TR3337_KE plasmid by QuikChange protocol [3] with the primer QuikChange_KW_Top and the primer QuikChange_KW_Bottom. Finally, the GUS region of the two plasmids above was amplified with the primer InFusion_GUS_Not_Back and the primer GUS-GSG-FLAG_Xho_For to get the GUS-TR3337_KE-FLAG and GUS-TR3337_KW-FLAG fragments. The fragments were inserted into the linearized pET32-VHH(Caf)-GS-GUS-TR3337_KE/KW vectors (digested by restriction enzymes NotI and XhoI) by In-Fusion Assembly (Takara Bio) to achieve the pET32-VHH-GS-GUS-TR3337_KW-FLAG and pET32-VHH-GS-GUS-TR3337_KE-FLAG for expressing the prototypes of the $GUS_{TR3}$-based immunosensors.

To introduce the additional diagonal interface mutation, the DNA fragment TR3-514A was amplified from the pET32-VHH-GS-GUS-TR3337_KW-FLAG with the primer InFusion_GUS_Not_Back and the primer BZ-GUSTR3337-KEKW-H514A-R. The DNA fragments TR3-KW-FLAG and TR3-KE-FLAG were amplified from plasmids pET32-VHH-GS-

GUS-TR3337_KW-FLAG and pET32-VHH-GS-GUS-TR3337_KE-FLAG with the primer BZ-GUSIV5KW-H514A-F/BZ-GUSIV5-TR-KE-H514A-F and the primer BZ-GUSFlag-XhoI-R, respectively. The fragment TR3-514A, TR3-KW-FLAG or TR3-KE-FLAG, and the linearized pET32-VHH-GS-GUS-TR3337_KW-FLAG (by NotI and XhoI) were ligated with In-Fusion assembly to yield plasmids pET32-VHH-GS-GUS-TR3337_AKW-FLAG and pET32-VHH-GS-GUS-TR3337_AKE-FLAG, respectively.

**Combinatorial library construction**

The library DNA fragments containing random mutations at H514, K516, and W517 positions were prepared by PCR from the template pET32-VHH-GS-GUS-TR3337_KW-FLAG with the mixed primer BZ-TR3337-Lib1-Ins-F, BZ-TR3337-lib1Y-R, BZ-TR3337-lib1W-R, and BZ-TR3337-lib1E-R at a molar ratio of 1:0.33:0.33:0.33. The vector fragment was obtained by an inverse PCR from the template pET32-VHH-GS-GUS-TR3337_KW-FLAG with primer BZ-TR3337-Lib1-Vec-F and primer BZ-TR3337-Lib1-Vec-R. The library DNA fragments and the vector fragment were ligated by using In-Fusion assembly and transformed into XL10-Gold competent cells. Over 10,000 transformants were pooled and cultivated for extracting the library plasmids.

**BSA-caffeine conjugation and ELISA**

The caffeine-BSA conjugate was prepared according to the published protocols [4,5] with modifications. Two mg of theophylline-7-acetic acid (Tokyo Chemical Industry) was dissolved in 450 µL of sodium acetate buffer (50 mM, pH 5.8) as a hapten solution. Ten mg of BSA was dissolved in 1 mL of ultra-pure water as a BSA solution. 450 µL of the hapten solution and 200 µL of the BSA solution were mixed then added into a new microtube containing 10 mg EDC. The reaction was mixed by pipetting immediately and incubated at room temperature for 2 h, then 4 °C overnight. The conjugate was then diluted with 10 mM phosphate-buffered saline (PBS) and glycerol to a final BSA concentration of 2 mg/mL with 30% glycerol, and stored at −30 °C.

The caffeine-BSA conjugate and BSA control were coated on a 96-well microplate at a concentration of 10 µg/mL. After incubation for 2 h at room temperature, the solution was removed and the microplate was blocked with 20% (v/v) ImmunoBlock (KAC, Hyogo, Japan) in PBST. After 1.5 h, the plates were washed three times with PBST (PBS with 0.1% Tween 20), and 100 nM $V_{HH}$-$GUS_{TR3}$-KE (in PBST containing 5% ImmunoBlock) was added into each well. After incubation for 1.5 h, the plate was washed three times with PBST, followed by the addition of 100 µL HRP-conjugated anti-6×His antibody (HRP-66005, Proteintech; 1:4000 dilution in PBST containing 5% ImmunoBlock). After 1 h of incubation and washing, 100 µL of substrate solution (0.2 mg/mL 3,3′,5,5′-Tetramethylbenzidine and 30 mM $H_2O_2$ in 100 mM sodium acetate buffer, pH 6.0) was added to develop the assay, and the absorbance at 450 nm and 655 nm was measured by a microplate reader SH-1000 (Corona Electric, Ibaraki, Japan) after terminating the reaction with 50 µL of 10% $H_2SO_4$.

#### Supplementary Figure

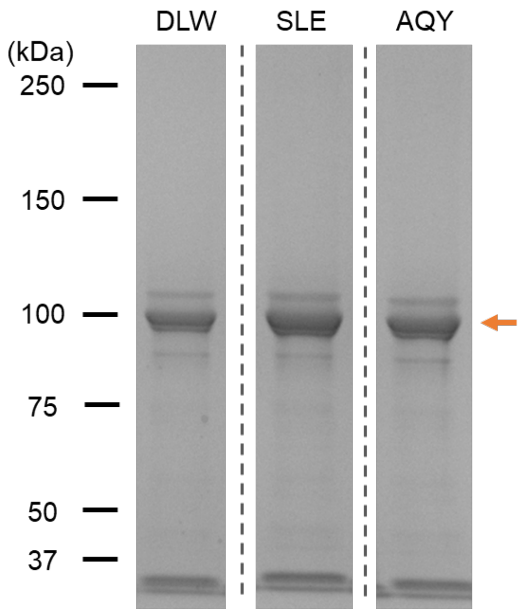


**Fig. S1. SDS-PAGE analysis of the purified immunosensor variants**. DLW: $V_{HH}$-$GUS_{TR3}$-DLW; SLE: $V_{HH}$-$GUS_{TR3}$-SLE; AQY: $V_{HH}$-$GUS_{TR3}$-AQY; Orange arrow indicates the monomer of the immunosensors. Dash lines divided the grouped lanes that are from the same or different gels.

#### Supplementary Tables

##### Table S1. Oligonucleotides and gene fragments used in this study.

| Name | Sequence |
|---|---|
| Q493R_top | 5′-CTTCTGGCCTGGCGGGAGAAACTGC-3′ |
| GUS_XhoFor | 5′-CCGCTCGAGTAGTCATTGTTTGCCTCCCTG-3′ |
| GUS_NotBack | 5′-ATAAGAATGCGGCCGCTATGTTACGTCCTGTAGAAA-3′ |
| Q493R_bottom | 5′-CCGCCAGGCCAGAAGTTCTT-3′ |
| QuikChange_KW_Top | 5′-CGGCCTGCACTCAAAGTGGACCGACATGTGGAGT-3′ |
| QuikChange_KW_Bottom | 5′-ACTCCACATGTCGGTCCACTTTGAGTGCAGGCCG-3′ |
| InFusion_GUS_Not_Back | 5′-GGCGGTAGCGCGGCCGCTATGTTACGTCCTGTAGAAA-3′; |
| GUS-gsg-FLAG_Xho_For | 5′-GGTGGTGGTGCTCGAGTCACTTGTCATCGTCGTCCTTGTAGTCACCGGA TCCTTGTTTGCCTCCCTGCTG-3′ |
| BZ-GUSTR3337-KEKW-H514A-R | 5′-CTTTGAGGCCAGGCCGGCTAACGCATCC-3′ |
| BZ-GUSIV5KW-H514A-F | 5′-GCCTGGCCTCAAAGTGGACCGACATGTGG-3′ |
| BZ-GUSIV5-TR-KE-H514A-F | 5′-GGCCTGGCCTCAAAGGAGACCGACATGTGG-3′ |
| BZ-GUSFlag-XhoI-R | 5′-GTGGTGGTGGTGCTCGAGTC-3′ |
| BZ-TR3337-Lib1-Ins-F | 5′-ATGTTACGTCCTGTAGAAACCCCAAC-3′ |
| BZ-TR3337-lib1Y-R | 5′-CACTCCACATGTCGGTATACWKTGASNNCAGGCCGGCTAACGCATC-3′ |
| BZ-TR3337-lib1W-R | 5′-CACTCCACATGTCGGTCCACWKTGASNNCAGGCCGGCTAACGCATC-3′ |
| BZ-TR3337-lib1E-R | 5′-CACTCCACATGTCGGTTTCCWKTGASNNCAGGCCGGCTAACGCATC-3′ |
| BZ-TR3337-Lib1-Vec-F | 5′-ACCGACATGTGGAGTGAAGAGTATC-3′ |
| BZ-TR3337-Lib1-Vec-R | 5′-TTCTACAGGACGTAACATAGCGG-3′ |
| Synthesized C-terminal sequence of the $GUS_{TR3}$ | 5′-TGGCGGGAGAAACTGCATCAGCCGATTATCATCACCGAATACGGCGTG GATGCGTTAGCCGGCCTGCACTCAAAGGAGACCGACATGTGGAGTGAAG AGTATCAGTGTGCATGGCTGGATACGTATCACCGCGTCTTTGATCGCGTC AGCGCCGTCGTCGGTGAACAGGTATGGTCTTTCGCCGATTTTGCGACCTC GCAATCCATATTGCGCGTTGGCGGTTCCAAGAAGGGGATCTTCACCCGCG ACCGCAAACCGAAGTCGGCGGCTTTTCTGCTGCAAAAACGCTGGACTGGC ATGAACTTCGGTGAAAAACCGCAGCAGGGAGGCAAACAA-3′ |

**Table S2. Diagonal interface residues of *E. coli* GUS.**

| Residues (Wild-type) | Secondary structure | Interaction with diagonal subunit | Substitution in IV5 | Substitution in TR3337 |
|---|---|---|---|---|
| H313 | α3 Helix | | | |
| A511 | Loop | | | |
| G512 | | | | |
| L513 | | | | |
| **H514** | | α10 Helix E523 | | |
| M516 | Loop | | | |
| Y517 | | | F | |
| E523 | α10 Helix | H514 | | |
| E524 | | | | |
| K576 | Loop | | | |
| P577 | | | | |
| S579 | α11 Helix | | | |
| F582 | | | Y | |

**Table S3. Alignment of the sequences with > 40% identity of *E. coli* GUS (residue 512 - 525).**

| Organism | UniProt ID | Identity | Sequence from 512-525 |
|---|---|---|---|
| *E. coli* K12 | P05804 | 1.000 | GLHSMYTDMWSEEY |
| *Shigella flexneri* | A0A0H2UZK0 | 0.993 | GLHSMYTDMWSEEY |
| *Salmonella enterica* | A0A4Z0PJH2 | 0.819 | GLHSMYSDMWSEEY |
| *Celerinatantimonas diazotrophica* | A0A4R1J8H9 | 0.645 | GLHSIYNEMWSEEY |
| *Saccharopolyspora coralli* | A0A5Q3QAS3 | 0.571 | GLHSVTPQPWSEEY |
| *Aspergillus parasiticus* (**Beta-Mannosidase**) | A0A5N6E663 | 0.549 | GLHSVMVTPWSEEF |
| *Microlunatus sp. KUDC0627* | A0A516Q1J0 | 0.506 | GMHQLIAQPWSEEY |
| *Bacillus cellulosilyticus* (**Beta-galactosidase**) | E6TZA4 | 0.481 | GFHQDPPVMFTEEY |
| *Homo sapiens* | P08236 | 0.454 | GFHQDPPLMFTEEY |
| *Mus musculus* | P12265 | 0.455 | GIHEDPPRMFSEEY |
| *Danio rerio* | F1QGS9 | 0.439 | GLHSDPPMMFTEEY |
